# Investigating olfactory behaviors in adult zebrafish

**DOI:** 10.1101/859033

**Authors:** Florence Kermen, Lea Darnet, Christoph Wiest, Fabrizio Palumbo, Jack Bechert, Ozge Uslu, Emre Yaksi

## Abstract

Odor-driven behaviors such as feeding, mating and predator avoidance are crucial for animal survival. While the zebrafish olfactory circuitry is well understood, a comprehensive description of odor-driven behaviors is needed to better relate olfactory computations to animal responses. Here, we used a medium-throughput setup to measure the swimming trajectories of 10 zebrafish in response to 17 ecologically relevant odors. By selecting appropriate locomotor metrics, we constructed ethograms systematically describing odor-induced changes in the swimming trajectory. We found that fish reacted to most odorants, using different behavioral programs and that combination of few relevant behavioral metrics enabled to capture most of the variance in these innate odor responses. We observed that monomolecular odors in similar chemical categories were weakly clustered based on the behavioral responses, likely because natural odors elicited stronger reactions than the monomolecular odors. Finally, we uncovered a previously undescribed intra and inter-individual variability of olfactory behaviors and suggest a small set of odors that elicit robust responses. In conclusion, our setup and results will be useful resources for future studies interested in characterizing olfactory responses in aquatic animals.

## INTRODUCTION

Olfactory cues are powerful drivers of a wide range of behavioral responses in fish, which are related to reproduction, foraging, fear and anxiety (Hara, 1986; Keller-Costa, Canário, & Hubbard, 2015; Florence Kermen, Franco, Wyatt, & Yaksi, 2013). The neural pathways underlying these stereotyped locomotor responses have been the focus of extensive research and are well described (Derjean et al., 2010; Fore, Cosacak, Verdugo, Kizil, & Yaksi, 2019; Friedrich & Korsching, 1998; Jetti, Vendrell-Llopis, & Yaksi, 2014a; Florence Kermen & Yaksi, 2019; Koide et al., 2009; Miyasaka et al., 2014; Wakisaka et al., 2017; Yabuki et al., 2016; Yaksi, Judkewitz, & Friedrich, 2007; Yaksi, von Saint Paul, Niessing, Bundschuh, & Friedrich, 2009). Odors strongly activate the reticulo-spinal cells controlling locomotion in the sea lamprey, via a circuit involving the olfactory bulb, posterior tuberculum and mesencephalic area (Derjean et al., 2010). Food-related odorants activate hypothalamic regions involved in appetite control in zebrafish (Wakisaka et al., 2017) and evoke foraging behavior in a wide range of fish species (Kaniganti et al., 2019; Koide et al., 2009; Lindsay & Vogt, 2004; Savoca, Tyson, McGill, & Slager, 2017; Wagner, Stroud, & Meckley, 2011; Wakisaka et al., 2017). Alarm cues activate regions involved in adaptive fear response that are homologous to the mammalian basolateral amygdala, septum and paraventricular nucleus of the hypothalamus (Faustino, Tacão-Monteiro, & Oliveira, 2017), and evoke anti-predatory behavior (Barreto et al., 2013a; Pfeiffer, 1963; v. Frisch, 1942; Zhao & Chivers, 2005). Thus, a precise characterization of the link between ecologically relevant olfactory cues and odor-driven behaviors is an important step towards characterizing the neural circuits generating these essential behaviors and how they are affected by animal’s internal states, such as satiety, fear or anxiety. Paradoxically, while the fish olfactory circuitry is well characterized, a comprehensive description of zebrafish behavior in response to ecologically relevant odors is needed to better relate olfactory computations to animal behavior.

A growing number of studies has begun to address this gap in knowledge by characterizing the change in zebrafish swimming patterns in response to olfactory cues, identifying clear negative (avoidance) and positive (approach) chemotactic responses (Braubach, Wood, Gadbois, Fine, & Croll, 2009; Faustino et al., 2017; Hinz et al., 2013; Hussain et al., 2013; Koide et al., 2009; Lindsay & Vogt, 2004; Mann, Turnell, Atema, & Gerlach, 2003; Mathuru et al., 2012; Vitebsky, Reyes, Sanderson, Michel, & Whitlock, 2005; Wakisaka et al., 2017; Yabuki et al., 2016). These studies gathered behavioral responses to cues belonging to one or two of the following odor categories: food-related cues (Koide et al., 2009; Wakisaka et al., 2017), social-related cues (Hinz et al., 2013; Yabuki et al., 2016), decay-related cues (Hussain et al., 2013), alarm cues (Faustino et al., 2017; Mathuru et al., 2012). This approach precludes the comparison of behavioral responses between odor categories in the same individual, which could be useful to uncover specific stereotyped motor programs. Moreover, olfactory behaviors were either measured in groups of fish (Faustino et al., 2017; Lindsay & Vogt, 2004; Yabuki et al., 2016), thus masking potential inter-individual variability in odor sensitivity or preferences, or the inter-individual variability was not specifically quantified (Hussain et al., 2013; Koide et al., 2009; Mathuru et al., 2012; Vitebsky et al., 2005; Wakisaka et al., 2017). Therefore, there is a need for testing behavioral responses of individual fish to a broad range of odorants spanning the natural stimulus space.

Here we characterize zebrafish olfactory behavior using a medium-throughput setup allowing for exposure to well-defined odor concentrations. Using this approach, the swimming trajectories of 10 fish were recorded in response to 17 ecologically relevant odors. By selecting 7 appropriate locomotor metrics, we constructed behavioral ethograms systematically describing odor-induced changes in the swimming trajectory. We found that fish reacted to most odorants, using different behavioral programs. A combination of few relevant behavioral metrics enabled to capture most of the variance in these innate odor responses. In general, odors belonging to similar categories were weakly clustered based on the behavioral responses. This was likely because natural odor extracts (food, blood, skin extract) have a tendency to elicit stronger reactions than the corresponding individual monomolecular components. Finally, we quantified intra and inter-individual variability of olfactory behaviors and suggest a small set of odors that elicit robust responses. In conclusion, both our setup and our results will be useful resources for future studies interested in characterizing olfactory responses in aquatic animals.

## RESULTS

### A vertical olfactory setup with precise control of olfactory cue concentration and fast switching of odors

To reproducibly measure fish responses to a large variety of odorants, we built a computer-controlled setup automatically recording the position of freely swimming individual fish (**Figure 1A**). The arena was 15 cm large, 11.5 cm high, and 3 cm deep (approximately 6 × 5 × 1 fish body lengths) and contained around 400 mL of water, allowing us to investigate zebrafish displacement in both the vertical and horizontal dimensions. The flow rate was adjusted to 90 mL/min, which was fast enough to rapidly clear the arena, but not strong enough to exhaust or stress the fish. In addition, a T-shaped connector deflected the inflow towards the lateral walls (**Figure 1A**). This was to avoid pushing the fish down and to enable a rapid and homogeneous distribution of the stimuli within the arena. To characterize the onset and dynamic of the olfactory cue delivered to the arena, we replaced it by a dye and measured the change in reflected light overtime (**Figure 1B**). The cue reached the arena 8 seconds after the valve opened and rapidly spread through the arena, covering its entire volume within 30 seconds. The cue concentration had returned to pre-stimulus levels within 15 minutes (**Figure 1C**). Based on this, we chose an inter-trial duration of 20 minutes to ensure complete clearance of prior cues before the following recording started.

**Figure 1:**
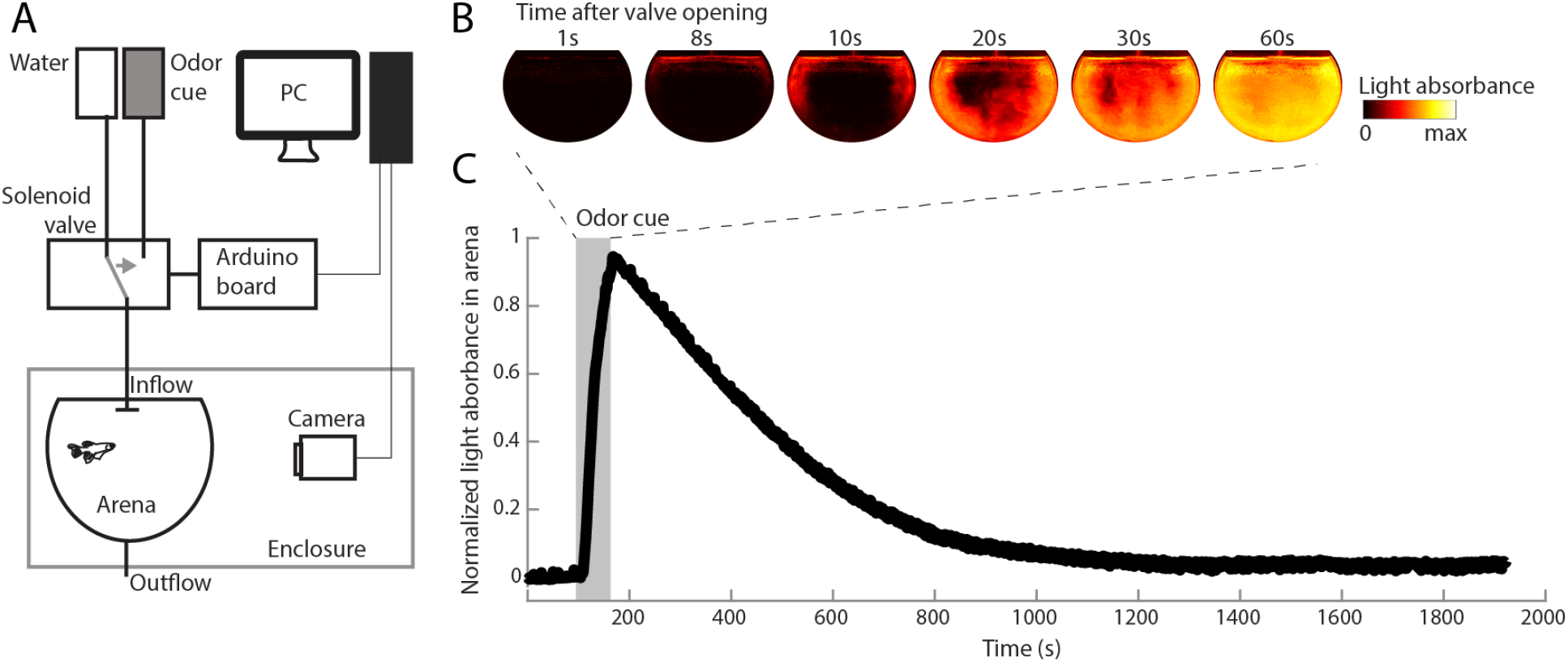
Experimental setup. **A)** Schematic of the experimental setup enabling to deliver olfactory cues to the arena where the fish is swimming using a computer-controlled solenoid valve. **B)** Image series showing the rapid and evenly spread of the olfactory cue within the arena after valve opening. The olfactory cue was replaced by a dye whose normalized concentration in the arena is plotted over time (0 = no stimulus; 1 =same concentration as in the odor cue bottle). **C)** Dynamic of stimulus concentration in the arena over the course of 30 minutes. The valve was open for 1 min.

### Characterization of zebrafish behavior in response to diverse olfactory cues

Fish rely on different categories of water-soluble olfactory cues to guide fundamental behaviors important for survival. We thus chose ecologically relevant olfactory cues related to one of the four following categories: feeding, social, decay and alarm cues (see **Table 1**). We used seven different feeding-related odorants: five amino-acids, which are substances eliciting foraging (Koide et al., 2009); a mix of nucleotides signaling the freshness of food (Wakisaka et al., 2017); and food cues extracted from fish-food flakes. Social-related odorants consisted of chemicals excreted by conspecifics: a mix of bile acids, which have been shown to guide sea lampreys to spawning sites (Peter W. Sorensen et al., 2005); ammonium and urea, which are metabolites present at high concentration in fish urine (Braun, Steele, Ekker, & Perry, 2009); and prostaglandin 2α, a fish reproductive pheromone released by ovulated females (Yabuki et al., 2016). Odor cues signaling decay consisted of putrescine, cadaverine and spermine, which are three amines enriched in decaying flesh and avoided by zebrafish (Hussain et al., 2013). Finally, alarm cues consisted of zebrafish skin extract (Døving & Lastein, 2009), zebrafish blood (Barreto et al., 2013b) and chondroitin sulfate, a compound previously identified as a component of fish skin extract that elicited similar alarm response (Mathuru et al., 2012).

**Table 1:** List of odorants used.

| short name | full name | category | concentration | CAS number |
| --- | --- | --- | --- | --- |
| chond | chondroitine sulfate | alarm substance | 1 µg/mL water | 9082-07-9 |
| skin | skin extract | alarm substance | 0.2 g/300 mL water | see methods |
| blood | blood | alarm substance | 1 µl/10 mL water | see methods |
| cad | cadaverine | decay odor | 10<sup>-4</sup> mol/L | 462-94-2 |
| put | putrescine | decay odor | 10<sup>-4</sup> mol/L | 33-93-7 |
| sperm | spermine | decay odor | 10<sup>-4</sup> mol/L | 71-44-3 |
| phe | L-Phenylalanine | foraging | 10<sup>-4</sup> mol/L | 63-91-2 |
| arg | L-arginine | foraging | 10<sup>-4</sup> mol/L | 74-79-3 |
| his | L-histidine | foraging | 10<sup>-4</sup> mol/L | 5934-29-2 |
| ala | L-alanine | foraging | 10<sup>-4</sup> mol/L | 56-41-7 |
| met | L-methionine | foraging | 10<sup>-4</sup> mol/L | 63-68-3 |
| cys | L-cysteine | foraging | 10<sup>-4</sup> mol/L | 52-90-4 |
| glut | L-glutamic acid | foraging | 10<sup>-4</sup> mol/L | 56-86-0 |
| food | food odor | foraging | add concentration | see methods |
| nucl | nucleotides (IMP & AMP) | foraging | 10<sup>-4</sup> mol/L | 84-21-9 ; 352195-40-5 |
| bile | taurocholic acid | social interaction | 10<sup>-4</sup> mol/L | 345909-26-4 |
| amo | ammonium chloride | social interaction | 10<sup>-4</sup> mol/L | 12125-02-9 |
| pgf2α | prostaglandin 2α | social interaction | 10<sup>-7</sup> mol/L | 38562-01-5 |
| urea | urea phosphate salt | social interaction | 10<sup>-4</sup> mol/L | 4861-19-2 |
| water | water control | control |  |  |

To characterize zebrafish olfactory behaviors, we then measured the swimming trajectory of 10 adult fish (7 males and 3 females) in response to these 17 odorants and to a water control. Single fish were habituated to the arena and olfactory cues were delivered after a 5 min baseline period. Mapping the fish position after the odor cue delivery yielded occupancy maps that differed markedly across odorants (**Figure 2 A,B,C,D**). In particular, feeding cues such as food extract, nucleotides and methionine induced exploration of the upper part of the arena (**Figure 2A**), where the odor cue was first delivered. In contrast, fish swam at the bottom of the tank in response to alarm cues such as blood and skin extract (**Figure 2C**). Overall, except increased activity closer to the lateral walls, the average occupancy map in response to feeding (**Figure 2F**) and social cues (**Figure 2G**) showed no clear differences compared to the water control (**Figure 2E**). This was likely due to the important inter-cue variability within these categories. Average occupancy maps in response to alarm cues (**Figure 2H**) revealed a consistent increase in bottom diving, that was also observable, although to a lesser extent, in response to decay cues (**Figure 2I**).

**Figure 2.**
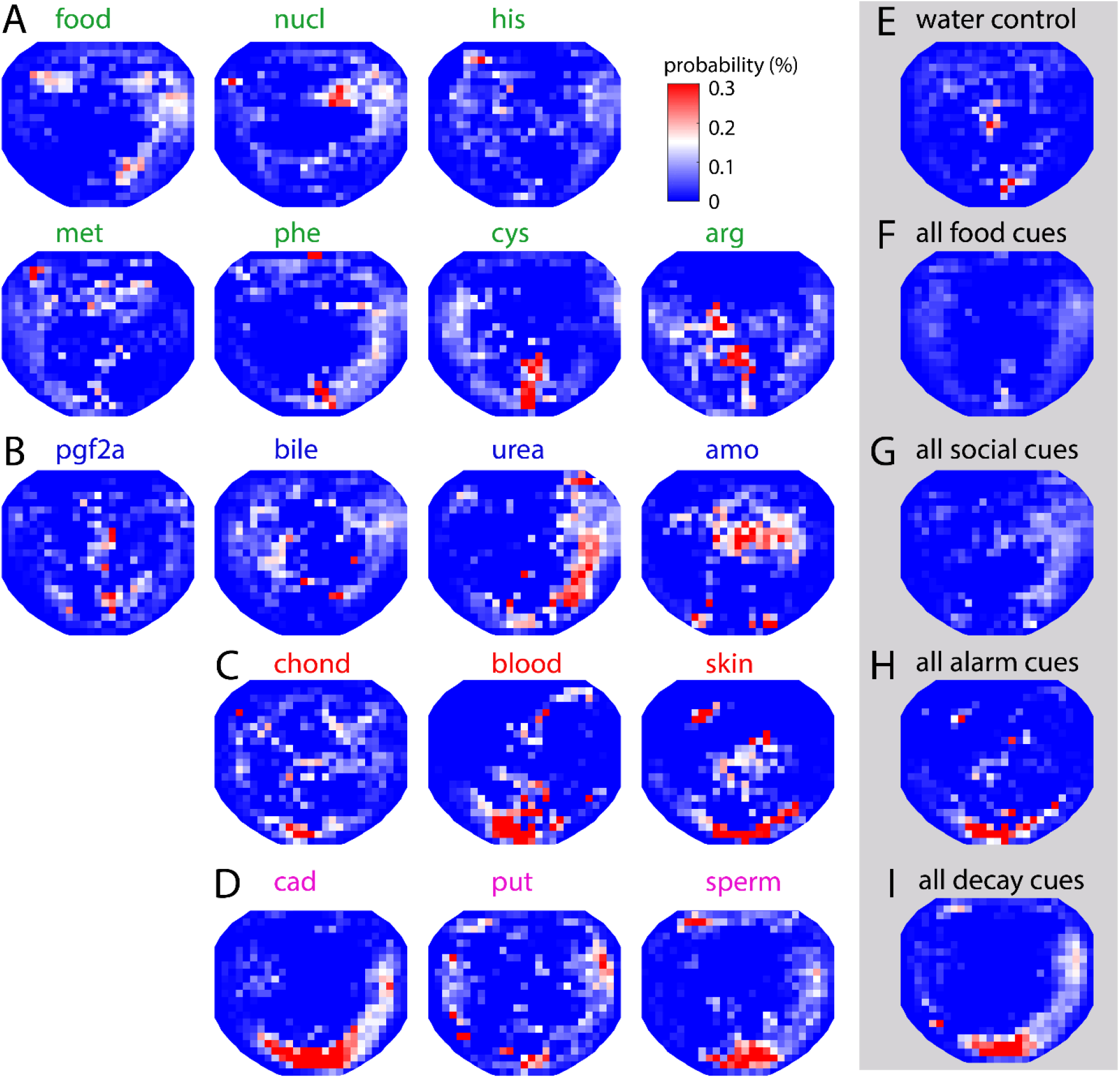
Spatial response of adult zebrafish to ecologically relevant odorants. Average occupancy maps representing the zones increasingly explored after onset of the olfactory cues (n=10 zebrafish): in response to **A**) Food-related cues (food extract, nucleotides, histidine, methionine, phenylalanine, cysteine and arginine, in green); **B**) Social-related cues (prostaglandin 2α, bile acids, urea and ammonium, in blue); **C**) Decay cues (putrescine, spermine, cadaverine, in magenta). **D**) Alarm cues (chondroitin sulfate, zebrafish blood, zebrafish skin extract, in red); **E**) Water control. **F**) Average of all food cues maps in A. **G**) Average of all social cues maps in B. **H**) Average of all alarm cues maps in C. **I**) Average of all decay cues maps in D.

### Quantification of odor-evoked changes in behavioral metrics

To quantify the dynamics of zebrafish locomotor behavior in response to odor cues, we calculated the odor-induced changes in the fish velocity (**Figure 3A**), the amount of burst swimming (**Figure 3B**) and the number of abrupt turns (**Figure 3C**), as well as the amount of horizontal (**Figure 3D**) and vertical swimming events (**Figure 3E**). To quantify the valence of odor cues, we also calculated the change in metrics reflecting decreased exploration, such as time spent freezing (**Figure 3F**) and the fish position along the vertical axis (**Figure 3G**), which are considered to be indications of anxiety and fear in fish. Representative examples of responses to selected olfactory cues are illustrated in **Sup. Figure 1**. Using these metrics, we built behavioral ethograms systematically describing the odor-induced changes in response to our diverse set of olfactory cues (**Figure 3A-G**). Importantly, we found no change in any of the behavioral metrics after water control delivery, indicating that small variation in flow rate or vibrations due to the valve opening and closing did not trigger behavioral responses.

**Figure 3:**
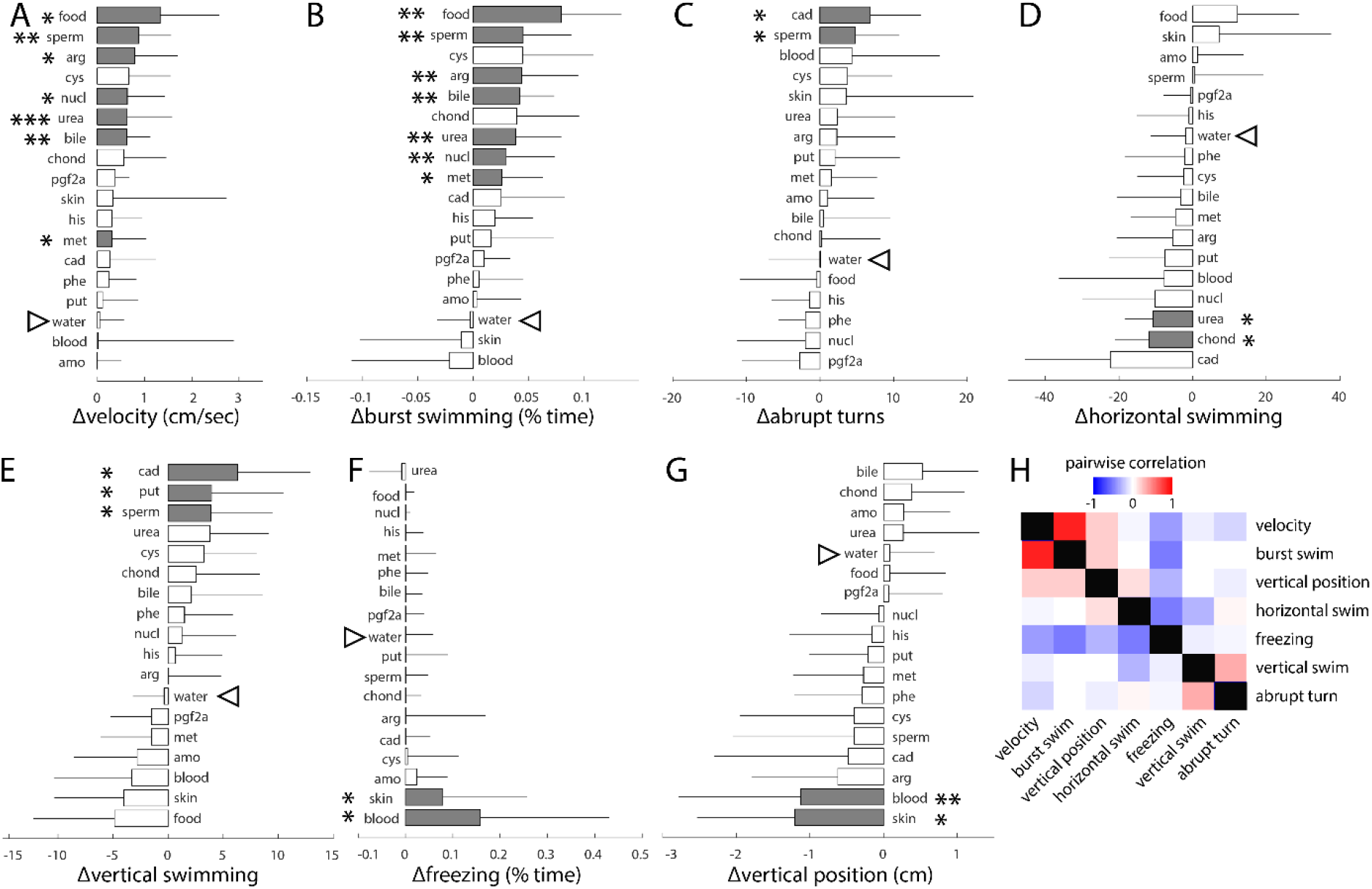
Behavioral responses of adult zebrafish to ecologically relevant odorants. Change in seven behavioral response metrics during the first two minutes following odor delivery: **A)** velocity, **B)** percentage of burst swimming, **C)** number of abrupt turns, **D)** horizontal swimming, **E)** vertical swimming, **F)** freezing, **G)** vertical position in the arena. The filled grey bars indicate significant differences from the water control, indicated by an arrowhead (Mann-Whitney U test; *p<0.05,**p<0.01 and ***p<0.001, see **Sup. Table 2** for all p values). Data are represented as mean + standard deviation. **H)** Pairwise correlation between all response indices in A-G.

Significant changes in swimming speed characterized by increased velocity and number of burst swimming events concerned the same odor cues: 4 food cues, 2 social cues and spermine. A significant increase in the amount of abrupt turns, possibly indicative of erratic swimming (Blaser & Gerlai, 2006), was observed in response to two decay cues (cadaverine and spermine). Interestingly, blood also elicited a marginally significant increase in sharp turns (**Sup. Table 1**, p=0.076). Changes in swimming strategy, in particular the amount of horizontal and vertical swimming have been observed in several fish species, related to foraging and spawning (Nakamura I, Watanabe YY, Papastamatiou YP, Sato K, & Meyer CG, 2011). Here, we observed a decrease in time spent swimming horizontally in response to urea and chondroitin. Interestingly, food extract elicited a marginally significant increase in the amount of horizontal swimming, a result reminiscent of foraging behaviors. Vertical swimming, up or down, was significantly increased in response to the 3 decay cues cadaverine, putrescine and spermine. Remarkably, the fear or anxiety-related behavioral indices freezing and vertical position in the tank, were significantly modulated solely after exposure to 2 out of 17 odors: the alarm cues skin and blood extracts. The time spent freezing and time spent lower in the arena increased in response to both of these cues, similar to what has been already been reported in response to skin extract (Blaser & Gerlai, 2006; Mathuru et al., 2012; Pfeiffer, 1963). Conspecific blood elicits the whole suite of specific alarm behaviors displayed in response to skin extract and thus seems to be a novel and equally powerful alarm substance in zebrafish.

To determine whether the behavioral metrics captured independent aspects of the odor response, we calculated their average pairwise correlation during odor response (**Figure 3H**). As could be expected, freezing was negatively correlated to most active locomotion indexes (velocity, burst swimming and horizontal swimming, **Figure 3H**), as well as to the vertical position in the tank, reflecting the fact that the majority of freezing took place at the bottom of the arena. To the exception of burst swimming that was strongly positively correlated to velocity, most metrics were weakly (anti-)correlated, indicating that they captured relatively independent aspects of the behavioral response.

### Consistency of odor responses within and between fish

The reproducibility or consistency of a behavior within and across individual is an important aspect of an animal’s response (Bell, Hankison, & Laskowski, 2009) quantifying how robust the observable behavior is over repeated presentation. Visual inspection of behavioral response metrics during an odor trial in our study indicated that fish could display similar responses to odor replicates (**Sup. Figure 1)**. For example, the patterns of freezing and vertical position in response to blood and the number of abrupt turns in response to skin extract are remarkably similar across replicates (**Sup. Figure 1A,C,G**). However, this is not the case for all odor cues, indicating that only a few specific odor cues might generate repeatable behaviors within fish. In order to quantify whether odor behavioral responses were repeatable, we measured the average correlation between behavioral responses to odor replicates (**Figure 4A**). Most odor cues elicited weakly correlated responses. However, cadaverine, blood, skin extract, food odor and bile acids displayed higher inter-replicate correlations ranging from 0.37 to 0.59 (**Figure 4A**), confirming that a specific subset of odors elicit consistent responses within fish. In order to determine whether odor cues elicited similar responses across fish, we calculated the average correlation in behavioral response between all pairs of fish for each odor (**Figure 4B**). Odors eliciting the most reproducible responses across fish were chondroitin sulfate, cadaverine, putrescine and urea, with between fish correlation ranging from 0.30 to 0.43. Interestingly, cadaverine elicited consistent responses both within and between fish (**Figure 4C**).

**Figure 4.**
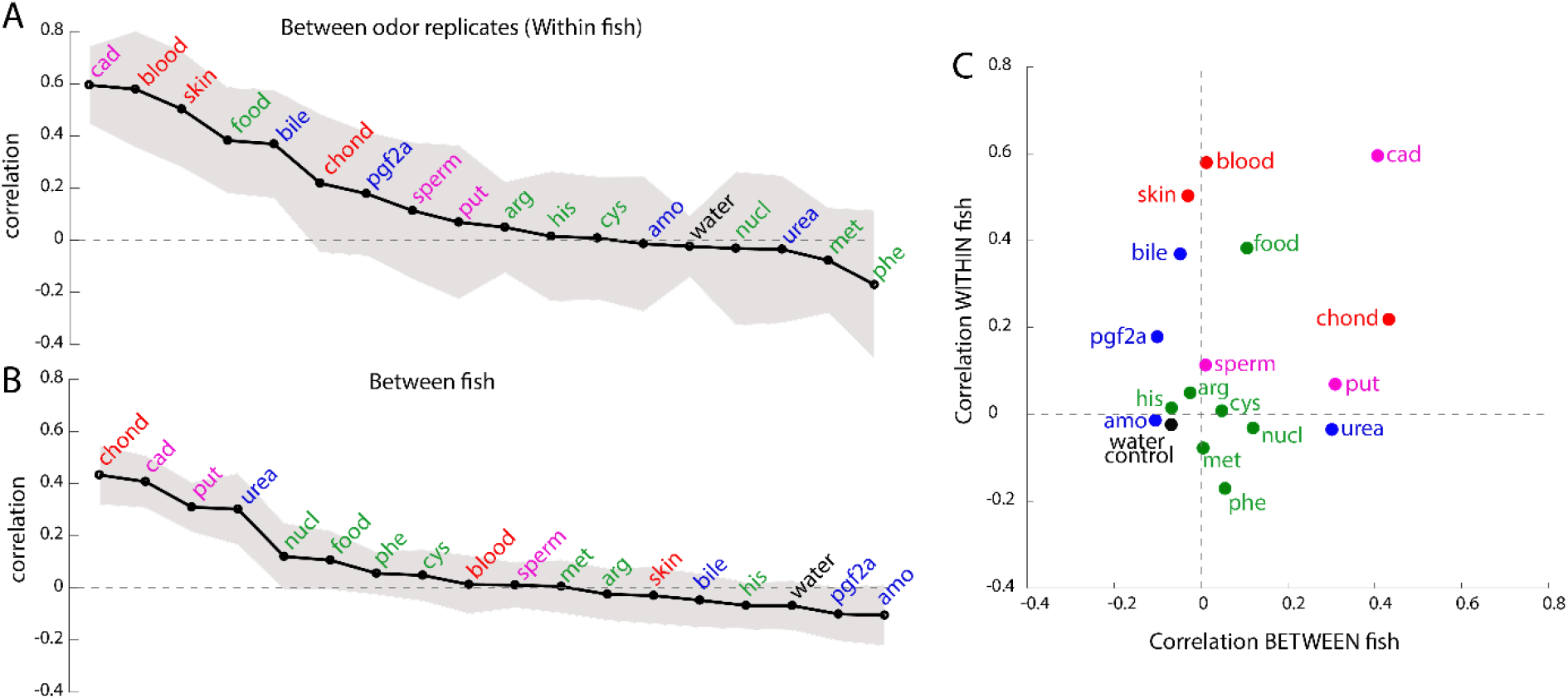
Consistency of odor responses within and between fish. **A)** Consistency of odor responses within individual fish. The average correlation (black line) between behavioral responses to replicates of the same odor within fish is plotted for all odor cues. Behavioral response was represented by a vector of the seven behavioral metrics described in Figure 3, averaged during the odor delivery period. **B)** Consistency of odor responses across different fish. The average correlation (black line) between the behavioral responses of all fish is plotted for all odor cues. The grey shaded area indicates s.e.m. **C)** Summary of data in A & B.

### Categorization of odors based on multi-dimensional olfactory behavior metrics

Next, we set to categorize odors based on behavioral responses, using a combination of all the metrics described above. To achieve this, we used hierarchical clustering to group olfactory cues eliciting similar behavioral responses (**Figure 5A**), based on the euclidean distances between all pairs of odors (**Figure 5B**). Clustering our broad set of odors based on zebrafish behavioral responses led to a categorization of odors. A first subset of amino acids, composed of methionine, histidine and phenylalanine, did not elicit responses prominently different from water control, likely due to the partial contribution of these monomolecular odors to feeding behavior. A second subset of amino acids, composed of cysteine and arginine, clustered together with the decay-related amine spermine. A third subset, composed solely of the natural feeding cue, food extract, elicited a different behavioral response from amino acids, which are monomolecular feeding cues. Similarly, although two natural alarm cues (blood and skin extracts) clustered together and were clearly different from all other odorants, the monomolecular alarm cue chondroitin sulfate was not part of that cluster.

**Figure 5:**
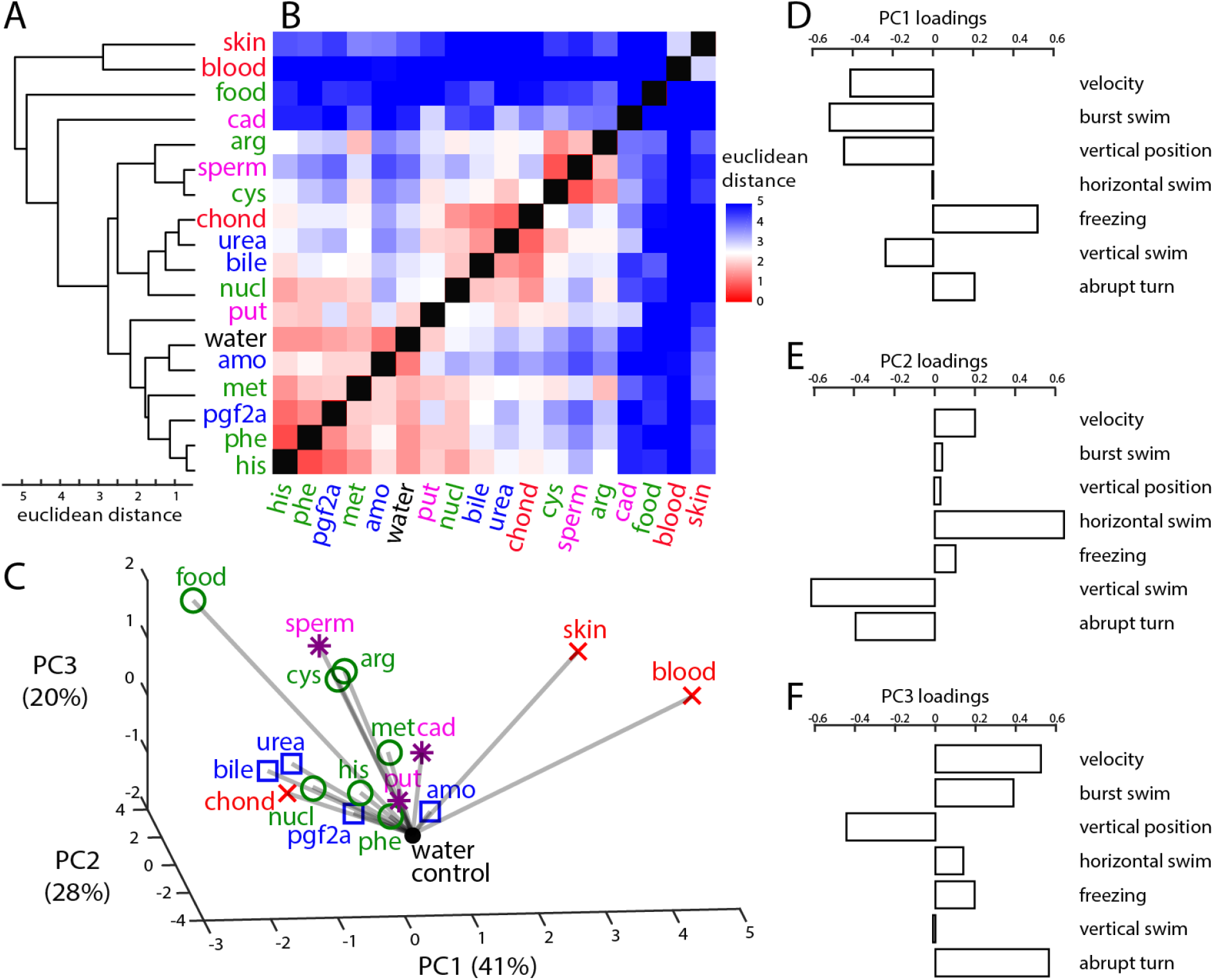
Behavior-based categorization of olfactory cues. **A)** Dendrogram representing the hierarchical clustering based on Euclidean distances between olfactory behaviors. **B)** Pairwise Euclidean distances between behavioral responses to all olfactory cues (7 response metrics, n=10 fish). Red indicates similar, and blue dissimilar, behavioral responses. **C)** Representation of all olfactory cues in the space composed of the first three principle components (PC) of the zscored median behavioral response across fish. The percentage of variance explained by each PC is indicated between brackets. Grey lines indicate the distance of each odor cue to the water control (filled black dot). Food-related cues (food extract, histidine, nucleotides, methionine, phenylalanine, cysteine and arginine) are represented by green circles. Social-related cues (bile acids, prostaglandin 2α, urea and ammonium) are represented by blue squares. Decay cues (putrescine, spermine, cadaverine) are represented by magenta asterisks. Alarm cues (chondroitin sulfate, zebrafish blood, zebrafish skin extract) are represented by red crosses. **D,E & F)** Respective contribution of all seven behavioral response metrics to PC1, PC2 and PC3.

To further facilitate the visualization of the complex behavioral response to the different odor categories, and to account for the dependencies between response measures (**Figure 3H**), we performed a dimensionality reduction analysis using principal component analysis (PCA, **Figure5C**). We found that the 3 first components of the behavioral space accounted for 89% of the variance, suggesting that olfactory responses can be explained by a combination of a few major behavioral programs.

The first PC clearly segregated food extract at one end and blood as well as skin extracts at the other end (**Figure 5C**; **Sup Figure 2A,C**). PC1 was positively correlated with decreased locomotion (velocity, burst swim), and increased fear or anxiety-like indices (bottom diving and freezing (**Figure 5D**)) thus possibly contrasting immediate rewards and threats. PC2 mostly separated animals movement in the horizontal versus vertical dimensions (**Figure 5E**), which is likely to represent evasive and exploratory behaviors. Finally, PC3 mostly represented parameters related to locomotion speed (**Figure 5F**).

Taken together, our results suggest that odors can be categorized based on olfactory behavioral metrics, yet the natural complex odors such as food, blood or skin extracts elicit clearly different and more prominent responses, when compared to monomolecular odors that were proposed to elicit stereotyped behaviors. We also observed that olfactory behaviors in response to 17 odors, can be explained by a handful of behavioral programs represented by the first three PCs.

## DISCUSSION

### Advantages and limitations of the medium-throughput olfactory behavior assay

We describe a medium-throughput setup to measure the behavior of individual fish exposed to olfactory cues of known concentrations. The automated delivery combined with rapid odor clearance enable to test multiple stimuli a day without manipulating the fish, while ensuring trial-to-trial reproducibility in stimulus dynamic. Precise control of cue concentration and dynamic is crucial for reproducible experiments, since it influences odor detection and valence (Semmelhack & Wang, 2009). Unlike studies in terrestrial animals where the delivery of airborne chemicals is controlled by olfactometers and measured by photoionization detectors (Johnson & Sobel, 2007; F. Kermen et al., 2011), measuring and rapidly clearing odors in the turbulent aquatic medium has proven challenging (Gerlai, 2011). Therefore, aquatic studies often confine the stimulus within the arena: in two‐current choice flumes (Hinz et al., 2013; Hussain et al., 2013), or using point source delivery (Koide et al., 2009; Mathuru et al., 2012; Wakisaka et al., 2017; Yabuki et al., 2016), which complicates the calculation of odor detection threshold and results in different odor exposure duration between fish. Here, we opted for local delivery combined with a rapid and homogeneous distribution of the cue. One drawback of our approach is to not allow for calculating a preference index (Hussain et al., 2013; Florence Kermen et al., 2016; Steck et al., 2012). However, we still observed clear responses to cues of opposite valence, in particular along the vertical dimension of the arena (**Figure 3**). In addition, a major advantage was to ensure spatially and temporally reproducible exposure over similar durations, a feature certainly useful for comparing inter-individual differences in olfactory thresholds.

### Metrics for fish locomotion

Locomotor activity of larval zebrafish is relatively well studied (Burgess & Granato, 2007; Mirat, Sternberg, Severi, & Wyart, 2013; Muto & Kawakami, 2013; Umeda & Shoji, 2017), however the metrics describing adult zebrafish behavior are scarce. Previous studies have used swimming speed and preference index to quantify olfactory behaviors. Zebrafish react to odor upon detection by an increase or a decrease in swimming speed. However, odors of different valence or ecological significance such as food related cues, and alarm cues can both increase swimming speed (Koide et al., 2009; Mathuru et al., 2012; Wakisaka et al., 2017). Moreover, the same fish can display biphasic responses to alarm substances, characterized by an alternation of freezing episodes (decreased speed) and bursts of erratic swimming (rapid disorganized swimming with sharp turning angles) (Gilson Volpato & Percília Giaquinto, 2001; Souza-Bastos, Freire, & Fernandes-de-Castilho, 2014). To best describe these temporally and spatially complex adult fish olfactory reactions, it is necessary to use a combination of metrics. Behaviors like erratic swimming are usually visually quantified by the experimenter (Døving & Lastein, 2009). Here, we used an entirely automated analysis to extract metrics classically used in the behavior analysis, such as velocity, burst of acceleration, vertical position in the tank, amount of freezing. We also used additional parameters such as amount of sharp turns > 90°, in order to quantify erratic movements and the patterns of vertical or horizontal swimming in order to examine the direction of swimming trajectory. Interestingly, we found that the 5 odors eliciting the largest increase in abrupt turns were 2 decay odors (cadaverine & spermine), 2 alarm cues (blood and skin extract) and an aversive amino acid (cysteine), indicating that the number of sharp turns could be an appropriate new metric to measure erratic movements in response to aversive stimuli.

### Responses to feeding cues

Fish detected most feeding cues as indicated by the significant increase in swim velocity and amount of burst swimming in response to 4 out of 7 food cues tested (nucleotides, methionine, arginine, and food extract). Testing each compound separately enabled us to uncover subsets of differing valence within amino acids. Indeed, cysteine and arginine clustered together with the decay-odor spermine and differed from other feeding cues in that they elicited more abrupt turns and bottom diving than other amino acids or the attractive nucleotides. This resonate with previous findings reporting that larval and adult zebrafish robustly avoid cysteine (Kaniganti et al., 2019; Vitebsky et al., 2005). Our results confirm this and propose arginine as an aversive amino acid.

### Responses to social cues

Bile acids are thought to be migratory cues guiding sea lampreys to spawning sites (Peter W. Sorensen et al., 2005). Here we found that they elicited an increased locomotion, similar to that evoked by feeding cues in this study and previous reports (Koide et al., 2009). Interestingly, urea evokes a very similar response to bile acid and was the closest related odorant in the behavioral space. No strong reaction was found in response to ammonium. Prostaglandin 2α is a steroid hormone produced by ovulated female fish that serves as a reproductive pheromone. It is detected by male fish olfactory system and contributes to the initiation of courtship in goldfish and zebrafish. Previous studies have shown that pgf2α exposure induces moderate behavioral response in male fish tested alone, but dramatic changes when tested in groups (P. W. Sorensen, Hara, Stacey, & Goetz, 1988; Yabuki et al., 2016). Here, we observed no noticeable response to pgf2α in individually tested fish, that were either males or females, confirming the necessity of group testing for this odor. This raises the interesting possibility that multimodal integration of olfactory information with additional sensory cues (vision, touch) is necessary to initiate attraction to pgf2α.

### Responses to alarm cues

Fish belonging to the Ostariophysi superorder (including zebrafish), display stereotyped anti-predator responses to chemical compounds released by damaged conspecific skin (Døving & Lastein, 2009). In agreement with this, we observed a significant increase in freezing and bottom diving in response to zebrafish skin extract. Blood can also be released when conspecifics are injured and was reported to decrease locomotion and increase latency to feed in the Nile Tilapia (Barreto et al., 2013a). Extending those findings, we show that low concentrations of conspecific blood (0.01%) elicited strong anti-predator responses (freezing and bottom diving) in adult zebrafish. Thus, our data suggest that anti-predator behavior in response to injured conspecifics within shoals could be jointly mediated by damage-released chemical cues present in the blood and skin.

### Responses to decay cues

Decomposition of fish flesh by bacteria releases ‘death-associated’ diamines such as cadaverine (Ben-Gigirey, Vieites Baaptista de Sousa, Villa, & Barros-Velazquez, 1999), which are avoided by adult zebrafish (Hussain et al., 2013). Interestingly, the decay odor spermine clustered with the aversive amino acid cysteine (Vitebsky et al., 2005). Both cadaverine and spermine produced significantly more abrupt turns than the water control, similar to responses to negative valence odorants (skin, blood, **Figure 3C**), suggesting that the decay cues were perceived as aversive. However, we found that decay odors induce only mild amounts of bottom diving and do not evoke freezing, which is consistent with previous reports (Hussain et al., 2013). To our knowledge, this is the first study that directly compares the responses to decay and alarm cues, which are released by dead or hurt animals. We found that decay cues did not cluster with the potent skin extract and blood alarm cues, confirming that they elicited a response that is qualitatively different from alarm cues in zebrafish. This finding is consistent with the ecology of these odors, given that alarm substances indicate a freshly wounded or killed fish, thus a high probability for an imminent threat, whereas bacteria-mediated production of decay cue takes hours to develop and thus signals a long-gone threat.

### Complex blends versus single compounds

Among feeding cues, the natural and complex food extract elicited the strongest response (**Figure 3A,B**), compared to simpler odors containing one or 2 monomolecular odorants, and was in general very different from all these cues (**Figure5A,B**). Similarly, skin and blood extracts evoked strong anti-predator responses, yet no such responses were observed to chondroitin sulfate in the same fish. It is unlikely that these differences are due to a lack of detection of the individual molecules, since the monomolecular odorants were presented at concentrations superior to the thresholds reported to elicit responses in zebrafish (Koide et al., 2009; Mathuru et al., 2012). Rather, this indicates that partial odor cues for feeding and alarm response do not elicit the full behavioral program.

### Inter- and intra-individual variability

Despite the increasing amount of studies interested in fish olfactory behaviors, few reports whether fish consistently respond to successive presentations of an odor cue (Imre, Di Rocco, Brown, & Johnson, 2016), and the robustness of odor response has not been systematically investigated for a broad range of odorants. We found that less than a third of the odors used here evoked reproducible responses within individual. Interestingly, cadaverine was the only monomolecular odor cue eliciting strongly consistent response both within and between fish. All other odors eliciting highly consistent responses within fish (bile acids, food, skin extract and blood), were only weakly correlated across fish. This could be due to the fact that individual zebrafish can use temporally consistent but different strategies for feeding (bottom vs top feeders) and defensive behaviors (proactive vs reactive coping styles) (Øverli et al., 2007). Conversely, chondroitin sulfate, urea and putrescine induced similar average responses across fish, but were only weakly consistent within fish. This emphasizes that intra- and inter-individual variability parameters describe different aspects of the behavioral responses. Thus, we suggest that odors with high intra-individual and low inter-individual reproducibility are useful stimuli to compare the behavioral and neural responses across fish with different personality traits, while investigating such individual differences.

In conclusion, our medium-throughput olfactory behavioral assay provides a low-cost and open setup to reproducibly measure olfactory reaction to multiple compounds in fish. Using this approach enabled us to collect an unprecedented amount of olfactory responses covering the natural stimulus space in adult zebrafish. We confirmed previously described behavioral responses to classically used odorants and also characterized a new powerful alarm substance (blood). We also provide recommendation for future studies to take into account the inter- and intra-individual reproducibility of odor behaviors. Finally, beyond neuroscience questions, olfaction psychophysics in fish also has important impacts in terms of species conservation. In this context, it is timely and crucial to provide tools and methods to reliably quantify the olfactory behavior in aquatic species to assess and ultimately limit the negative anthropogenic impact on aquatic ecosystems.

## METHODS

### Animal and housing

To allow future comparison with functional imaging studies, the behavioral experiment were carried out in transgenic zebrafish *Danio rerio* expressing the fluorescent calcium indicator Gcamp5 pan-neuronally (elavl3:GCaMP5 nacre,(Ahrens, Orger, Robson, Li, & Keller, 2013)). A total of 10 fish aged 6-12 months, including 3 females, were used. Fish measured on average 2.5 cm (+/− 0.16 std) from the tip of the nose to the base of the tail. They were kept in 3.5 liter tanks in a recirculating fish housing system (ZebTec Active Blue Stand Alone, Techniplast) at 28 °C under a 14:10 hour light/dark cycle. Two weeks before the start of experiments, pairs of fish were placed in a 3 L tank and isolated from each other by a transparent separator to keep track of individual fish day after day. Fish were fed once in the evening during the behavioral testing period and were thus food-deprived for 18h before testing. All animal procedures were performed in accordance with the animal care guidelines and approved by the Ethical Committee of KULeuven in Belgium and the Norwegian Food Safety Authority.

### Olfactory cues

Seventeen odorants, previously documented to activate the olfactory system in a range of aquatic species, and a water control were tested in this study. Single compounds odorants were purchased from Sigma Aldrich (Table1). Stock solutions were prepared, kept at −20°C, and diluted to final concentration in artificial fish water (AFW; 0.2 g/L marine salt in reverse osmosis water) the morning of the experiment. The final concentration to which the fish were exposed is documented in Table 1 for each olfactory cue. Food odor, blood, chondroitin sulfate and skin extract were freshly prepared the morning of the experiment. Food odor was prepared by incubating 1 g of commercially available fish food (SDS100, Scientific Fish Food) in 50 mL of AFW for 30 min. The solution was then filtered and further diluted 1:10 in AFW. For blood odor collection, adult nacre fish were rapidly euthanized in ice-cold water, the tail was cut close to the anal orifice and blood was collected from the dorsal aorta using a 20 µL pipette (Pedroso et al., 2012). 30 µL of blood was diluted in 300 mL of ice-cold AFW, filtered, and kept at 4°C until 1 hour before the start of the experiment the same day. Importantly, we made sure to sample blood from the dorsal aorta without touching the skin to avoid contamination by alarm cues released by epidermal club cells. For skin extract, adult nacre fish were rapidly euthanized in ice-cold water, decapitated and the skin was peeled off from the body. The collected skin (0.2 g) was incubated in 1 ml of AFW, mixed and centrifuged at 1300rpm and 4°C for 1h. 1 mL of the supernatant was then dissolved in 300 mL of AFW (Jetti, Vendrell-Llopis, & Yaksi, 2014b). Chondroitin sulfate was diluted the morning of the experiment as previously described (Mathuru et al., 2012), to avoid damage of this heavy molecule by freezing the stock solutions.

### Behavioral setup

Each fish was allowed to freely swim in a transparent semi-circular vertical arena (maximum depth= 15 cm; height= 11.5 cm; width: 3 cm) made of two transparent petri dishes (Falcon, 15cm diameter) glued together with epoxy. An opening was made at the top and the arena was equipped with inlet and outlet tubes. Water and odor stimuli were kept in glass bottles positioned on an elevated platform, providing a continuous inflow (90 mL/min), which was measured and adjusted using a flowmeter (Cole Parmer, 65mm flowmeter with a 3/16‘’ carboloy float). An outlet tube overflow ensured that the water volume contained in the arena remained constant. To avoid contamination between consecutive stimuli, the tubes were composed of Teflon (Cole Parmer, internal diameter: 3.2 mm). The arena was placed within a light-tight enclosure (black hardboard, Thorlab) to isolate the fish from visual interference, and a white LED covered by a diffuser provided homogeneous lighting inside the enclosure. Fish movement was recorded at 10 Hz using a camera (Manta223B, Allied Vision) positioned in front of the arena. Odorant delivery was automatically triggered in synchronization with video acquisition using a Matlab/C++ code and a simple electronic circuit composed of an Arduino Uno (Arduino) and a three-way solenoid valve (Biochem Fluidics). The dynamic of odorant stimulus in the arena was measured using a dye (methylene blue). The day before the experiment started, the fish was habituated to the arena for 3 hours, without odorant exposure. Then, at the beginning of each recording day, fish were allowed to habituate to the arena for at least 45 min before initiating recordings. Each trial started by 5 minutes without odorant, then the valve was switched open for 1 min allowing the odor stimulus to be delivered in the arena. Odors were delivered in random order. Zebrafish skin extract and zebrafish blood were delivered at the end of the recording session and no additional recordings were taken after these, due to their potent aversive behavioral effect. Successive trials were separated by at least 20 min, to allow for complete rinsing of odorant before the next trial started. An average number of 2.0 +/− 0.2 replicates per stimulus was collected. The experiments were conducted between 1-7 pm in a temperature controlled room (27 °C). The fish was returned to its home tank and fed every day at the end of the experiment.

### Swim trajectory extraction

The fish position (centre of mass) was automatically detected after background subtraction in Matlab (R2016a). Briefly, a background image was obtained by averaging all frames belonging to a given trial. The background was then subtracted from each individual frame. The background subtracted video was then processed by an erosion/dilation function. In rare cases where the corner of the arena or a moving drop of water was detected instead of the fish, a mask was applied to the image to constrain the detection within the arena, and the position was re-extracted. To correct for small day-to-day variation in camera’s position compared to the arena, the fish’s position was converted in cm and normalized across fish by setting the origin to the bottom left corner of the arena.

### Behavioral response metrics

We first calculated 7 metrics characterizing zebrafish behavior. Instantaneous speed (cm/sec) was calculated as the distance between successive positions divided by the recording frequency. Acceleration (cm^2^/sec) was the first derivative of speed. Freezing episodes were defined as immobility periods (speed<0.4 cm/sec) that lasted more than 5 seconds. The number of sudden swimming bursts (acceleration > 1cm^2^/sec) and the amount of sharp changes/turns in swim trajectory (turning angle > 90°) were also quantified. Horizontal and vertical swimming episodes were defined as number of events during which the fish swam with very little deviations (+/− 10°) from the horizontal and vertical lines, respectively. We then calculated a response metric associated with each behavioral parameter: difference between average value during 2 min preceding and following the stimulus onset. Positive values indicate an increase of the behavioral metric after stimulus delivery, whereas negative values indicate a decrease. Because we observed biphasic panic responses to alarm cues in most fish (escape first, then freezing), a longer post-odor time window lasting 4 min was used for quantifying freezing. These individual metrics were then averaged across replicates of the same odorants to yield a response matrix consisting of 7 metrics × 18 stimuli × 10 fish.

### Average occupancy maps

For each recordings, the arena was tilled into 5 mm squares and the percentage occupancy of each square was calculated based on the fish position. A differential occupancy map was then calculated for each odorant by subtracting the occupancy maps before and after odor delivery (during 4 min each). These differential occupancy maps were then averaged per odor and across fish to yield the average maps of areas increasingly explored after odor delivery shown in Figure 2.

### Statistical analysis

Data were examined for normality of distribution using a Shapiro-Wilk test in Matlab. As none of the metrics described here were normally distributed, difference from the water control condition were detected using a Mann Whitney U test, using a p value of 0.05 as threshold for significance.

## SUPPLEMENTARY FIGURES & TABLES

**Sup figure 1:**
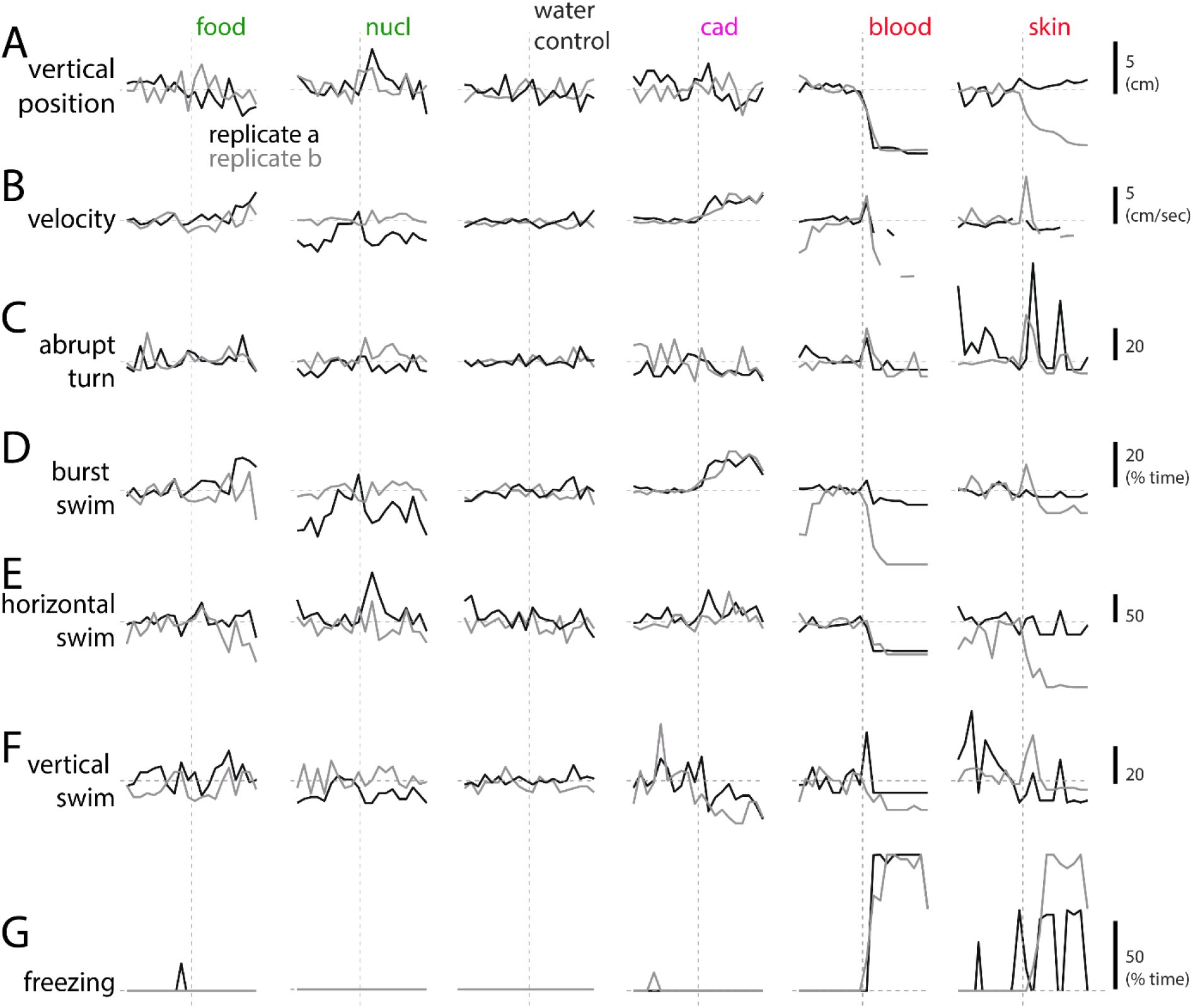
Example traces of behavioral parameters change in response to selected odor cues (from left to right : food odor, nucleotides, water control, cadaverine, blood and skin extract). All data come from the same fish. Replicates are indicates in black (replicate a) and grey (replicate b). Each behavioral parameters was averaged per time bins of 30 sec and normalized with respect to the baseline period (2 min before odor delivery). A to G: vertical position, Horizontal grey dotted line indicate 0. Vertical grey dotted lines indicate odor onset on each graph. The respective scales are indicated to the right for each parameters.

**Sup table 1:**
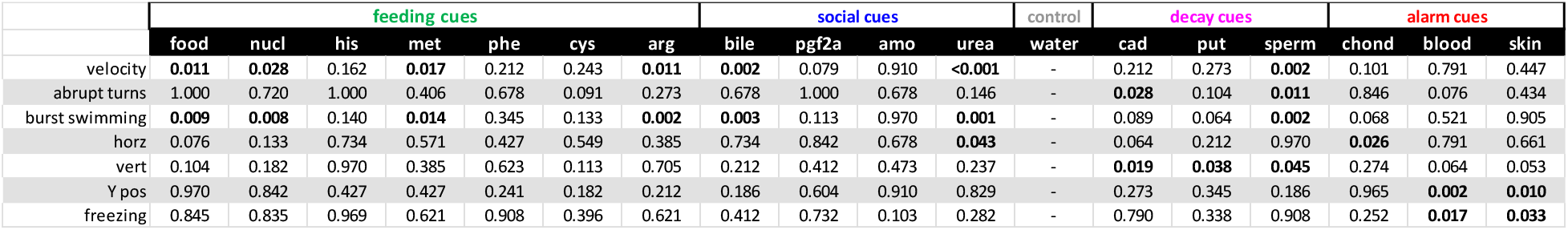
Outcome of statistical tests performed in Figure 3. **A-G.** P values corresponding to all odor cue – water control pairwise comparisons are displayed here. P values <0.05 are displayed in bold.

**Sup Figure 2:**
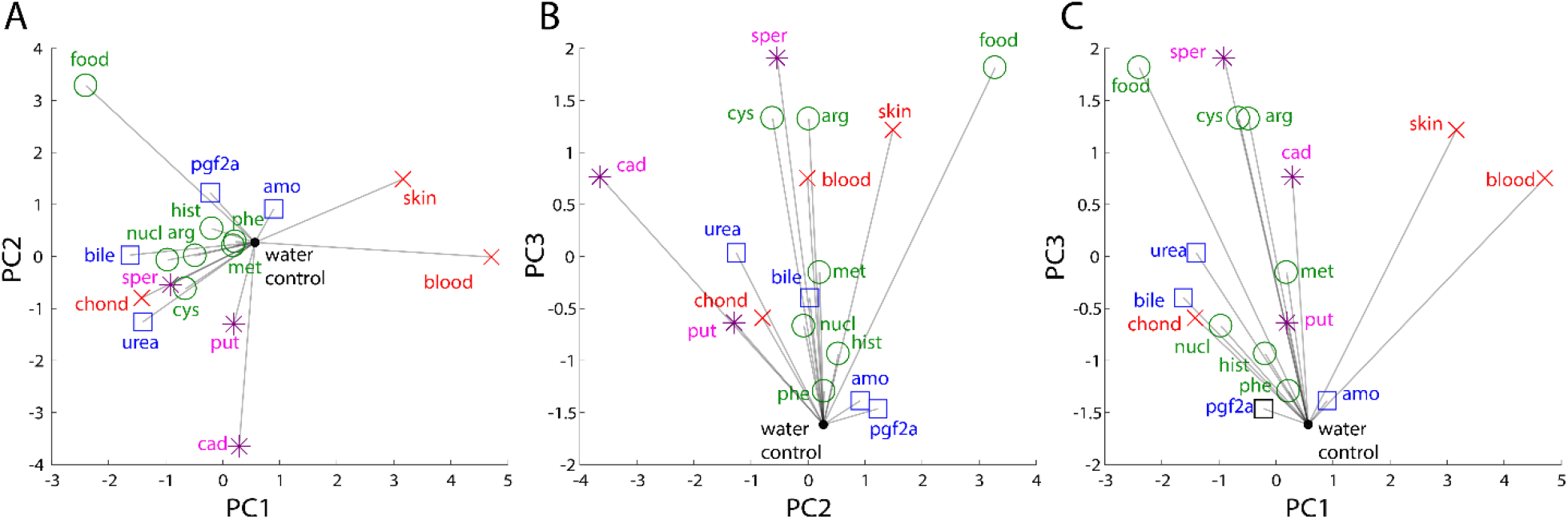
Representation of olfactory cues in the bidimensional spaces composed of behavioral PC1 & PC2 (A), PC2 & PC3 (B) and PC1 & PC3 (C). Grey lines indicate the distance of each odor cue to the water control (filled black dot). Food-related cues (food extract, histidine, nucleotides, methionine, phenylalanine, cysteine and arginine) are represented by green circles. Social-related cues (bile acids, prostaglandin 2α, urea and ammonium) are represented by blue squares. Decay cues (putrescine, spermine, cadaverine) are represented by magenta asterisks. Alarm cues (chondroitin sulfate, zebrafish blood, zebrafish skin extract) are represented by red crosses.

## CONTRIBUTION

EY & FK designed the study. FK, LD and CW built the experimental setup. FK, JB, FP and EY developed the tracking code. LD, CW collected the data with contribution from OU. FK analyzed the data and made the figures. FK and EY wrote the manuscript with inputs from all authors. EY conceived the study, supervised and trained the team members.

## FUNDING

This work was supported by NERF & V.I.B. at KU Leuven and the Kavli Institute for Systems Neuroscience at NTNU, as well as a Fyssen foundation post-doctoral fellowship (F.K.), a Boehringer Ingelheim fellowship (C.W.), ERC Starting Grant 335561 (E.Y.), a Norwegian Research Council FRIPRO grant 239973 (E.Y.) and a FRIPRO grant 262698 (F.K.).

